# rdacca.hp: an R package for generalizing hierarchical and variation partitioning in multiple regression and canonical analysis

**DOI:** 10.1101/2021.03.09.434308

**Authors:** Jiangshan Lai, Yi Zou, Jinlong Zhang, Pedro Peres-Neto

## Abstract

1. Canonical analysis, a generalization of multiple regression to multiple response variables, is widely used in ecology. Because these models often involve large amounts of parameters (one slope per response per predictor), they pose challenges to model interpretation. Currently, multi-response canonical analysis is constrained by two major challenges. Firstly, we lack quantitative frameworks for estimating the overall importance of single predictors. Secondly, although the commonly used variation partitioning framework to estimate the importance of groups of multiple predictors can be used to estimate the importance of single predictors, it is currently computationally constrained to a maximum of four predictor matrices.
2. We established that commonality analysis and hierarchical partitioning, widely used for both estimating predictor importance and improving the interpretation of single-response regression models, are related and complementary frameworks that can be expanded for the analysis of multiple-response models.
3. In this application, we aim at: a) demonstrating the mathematical links between commonality analysis, variation and hierarchical partitioning; b) generalizing these frameworks to allow the analysis of any number of responses, predictor variables or groups of predictor variables in the case of variation partitioning; and c) introducing and demonstrating the usage of the R package `rdacca.hp` that implements these generalized frameworks.

## Introduction

Canonical analysis (also called “constrained ordination”) considering multiple response variables (e.g., species) and multiple predictors (e.g., environmental features, spatial predictors) are widely used as inferential frameworks to determine and contrast the importance of multiple drivers (e.g., environmental conditions, traits) underlying the structure of ecological communities. Redundancy analysis (RDA; Rao 1964), canonical correspondence analysis (CCA; ter Braak 1986), and distance-based redundancy analysis (db-RDA; Legendre & Anderson 1999) are the most commonly used ones (Legendre & Legendre 2012). One central challenge in canonical analyses is the estimation of predictor contribution given that the number of regression parameters increase as a function of the number of response variables. For instance, if 200 response variables and 20 predictors are considered, one ends up with 4000 regression slopes.

Because canonical analyses are extensions of multiple regression models to multiple response variables (Peres-Neto *et al*. 2006), we can adapt the existing machinery of multiple regression models to tackle this challenge. The approach we develop here is based on generalizing commonality analysis, a single-response regression framework, to canonical analysis (i.e., multi-response variables). Commonality analysis is often used in psychology and education (Newton & Spurell 1967; Nimon & Reio 2011; see Ray-Mukherjee *et al*. 2014 for a rare application in ecology) to estimate the relative predictor contributions to the total model’s coefficient of determination (*R*^2^). Commonality analysis decomposes the total models’ *R*^2^ into unique fractions attributable to individual predictor and the shared fractions among predictors (covariation in the predictive space which can also identify multicollinearity issues). These fractions are semi-partial *R*^2^s (Peres-Neto *et al*. 2006) and allow circumventing well-known issues underlying variable importance and model interpretation based solely on standardized partial slopes (see Ray-Mukherjee *et al*. 2014 for a review of the issues). This is an interesting property because canonical analyses produce estimates of total models’ *R^2^* and semi-partial *R*^2^s that are unbiased (Peres-Neto *et al*. 2006).

Although commonality analysis is applied to single-response regression models, ecologists are widely familiar with the parallel framework of variation partitioning applied to canonical, with thousands of studies published using it (Borcard, Legendre & Drapeau 1992; Peres-Neto *et al*. 2006; see Fig. 1). Variation partitioning is employed by grouping predictors together into matrices and estimating the unique and shared semi-partial *R*^2^s of each matrix compounded across all response variables (Peres-Neto *et al*. 2006). Indeed, the algebra involved in commonality analysis and variation partitioning are equivalent even though they are often used in different situations. Notwithstanding, packages conducting commonality analysis are restrained to single response variables, e.g., R package `yhat` (Nimon, Oswald & Roberts 2013); and packages conducting variation partitioning (i.e., multi-response), such as the widely used R package `vegan` (Oksanen *et al*. 2006), are restrained to a maximum of four predictor matrices. These constraints are either implementational or computational. In particular, variation partitioning has not been generalized to multiple predictor matrices (but see Økland 2003) and commonality analysis has not been generalized to multi-response models.

**Figure 1.**
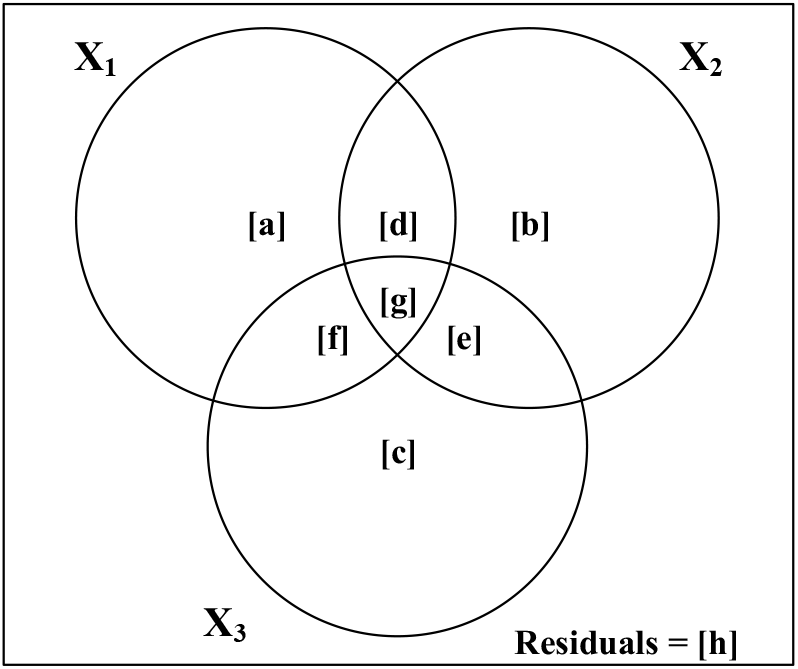
Venn diagram representing the variation partitioning of a response matrix **Y** regressed against three correlated predictors (or groups of predictors as in variation partitioning). All fractions add to 100% and the bounding rectangle represents the total variation in the response data (i.e., 100%) while each circle represents the relative portion of variation accounted by different fractions (see text for a detailed calculations and further explanation).

Beyond variation partitioning, estimating the relative importance of predictor in multiple regression (e.g., multicollinearity being an extreme case) is an old and very active research (see the reviews in Bi 2012; Nathans, Oswald & Nimon 2012; Grömping 2015). Among them, the “averaging over orderings” approach proposed independently by Lindeman, Merenda & Gold (1980), Cox (1985) and Kruskal (1987) in the 1980s has been considered as a breakthrough (Bi 2012). These methods are generally referred as to the LMG metric and are based on all model subsets, and are also equal to methods described independently such as hierarchical partitioning (Chevan & Sutherland 1991) and dominance analysis (Budescu 1993). Because the calculations described in these papers are overly complicated, one is led to think that they differ. In the next section, we simplify the presentation of “averaging over orderings” so that others can take full advantage of its simplicity while describing model complexity. Our computational presentation also makes a clear link between commonality analysis, and variation and hierarchical partitioning. Because we were particularly inspired by the very popular paper by Chevan & Sutherland (1991), we used the term “hierarchical partitioning” (HP) to relate to these equivalent methods. HP produces all possible combinations of predictors to determine the order in which a predictor dominates over the others (hence the name dominance analysis used by Badescu 1993) across all subset models, and it has been widely used and recommended to access the relative importance of predictors in multiple regression (Soofi 1992; Mac Nally & Walsh 2004; Walsh *et al*. 2004; Grömping 2006; 2007; 2009; 2015). Corresponding R packages are `relaimpo` for LMG (Grömping 2006), `hier.part` for HP (Walsh & Mac Nally 2013) and `dominanceanalysis` (Navarrete & Soares 2020). We show that analytical results from these packages are identical for multiple regression (see Appendix S1). However, unlike variation partitioning, HP is not currently available for multi-response models (i.e., canonical analysis) and is implemented in our package: `rdacca.hp`. By doing so, we also extend HP to consider matrices of predictors in variation partitioning.

The overall goal of this application paper is to unify commonality analysis, variation and hierarchical partitioning, generalizing them to unlimited number of responses and predictor variables (or matrices of predictors as in variation partitioning). These analyses are implemented in our package `rdacca.hp` which is presented and illustrated in the next sections.

## Unifying commonality analysis, and variation and hierarchical partitioning

Assuming a multi-response matrix **Y** and three correlated predictors (Fig. 1). The fractions of variation [a], [b] and [c] correspond to the variation in **Y** that are uniquely explained by predictors X_1_, X_2_, X_3_, respectively (i.e., unique semi-partial *R*^2^). Fractions [d], [e], [f] are the shared semi-partial *R*^2^s by combinations of their two respective predictors accounting for the third, Fraction [g] is the shared semi-partial *R*^2^ among all predictors, and [h] is the fraction corresponding to residuals. All fractions summed totalize 100%. Shared fractions are the variation in the response data that is explained by the correlation of the predictors involved. The larger this fraction is, the more multicollinearity is present in the model. While model selection reduces the shared variation among predictors (collinearity), it also reduces our ability to improve model interpretability. The calculations involved in variation partitioning are well described elsewhere (e.g., Peres-Neto *et al*. 2006) and our package: `rdacca.hp` generalizes it for any number of predictors (or matrices of predictors).

Individual predictor contribution in HP (also called “independent contribution”) can be simply estimated as its unique contribution to the total model *R*^2^ plus its average shared contributions with the other predictors, simplifying its presentation dramatically. The independent contribution across all possible models derived from combinations of predictors can be derived directly from variation partitioning (Fig. 1). For example, the independent contribution of X_1_ (*I*_*X*_1__) can be calculated by its unique contribution [a] and its respective shared contributions by the number of predictors involved in each of:

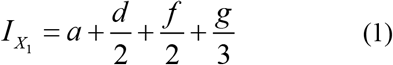

For completion, the contributions of X_2_ and X_3_ are calculated as follows:

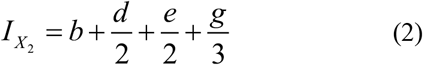

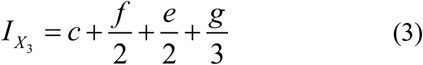

and:

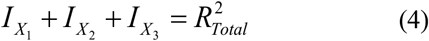

Generalizing, the independent contribution of any predictor *i* (*I*_*X*_*i*__) can be computed as:

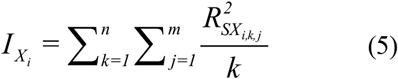

where *n* is the number of predictors, 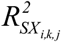 is semi-partial R^2^ of the *j^th^* fraction shared between X_i_ and the other *k* predictors, and *m* is the number of combinations that X_i_ shared with other *k* predictors, with 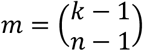. Note that as the number of predictor increases, the number of fractions including shared and unique *R*^2^s increases exponentially (2^N^-1 fractions). The above computation only uses individual predictors to illustrate the process. It is also applicable to matrices of predictors as in routine ecological applications applying variation partitioning.

In general, we find variation partitioning should be the starting point prior to hierarchical partitioning. While the former emphasizes unique and common variation among predictors, the latter emphasizes the overall importance of each predictor (or group of predictors). Our package `rdacca.hp` synchronously implements variation and hierarchical partitioning for single- and multiple-response models without limits in the number of predictors / matrices of predictors

## Package description

The `rdacca.hp` package is written in R (R Development Core Team 2019) and can be installed from CRAN (http://cran.r-project.org/web/packages/rdacca.hp/) or Github (https://github.com/laijiangshan/rdacca.hp). The package contains one key homonymous function: `rdacca.hp` that conducts both variation and hierarchical partitioning in single- and multiple-response multiple regression (canonical analysis). The internal function: `Canonical.Rsq`, calculates *R*^2^ and adjusted *R*^2^ (hereafter R^2^adj) of RDA, db-RDA and CCA, which are called by `rdacca.hp`. For canonical analysis, the R^*2*^adj is used given that the contribution of null predictors can differ quite a lot from zero due to sampling variation related to large number of predictors and small number of observations (Peres-Neto *et al*. 2006). The R^2^adj is calculated using Ezekiel’s formula (Ezekiel 1930) for RDA and db-RDA, while permutation procedure be used for CCA (Peres-Neto *et al*. 2006). The interpretation of arguments and some key notes in function `rdacca.hp` are described briefly here.

`rdacca.hp(dv, iv, method, type, n.perm, trace, plot.perc)`

In this usage, both a variation partitioning and hierarchical partitioning are performed in which the unique, shared (referred as to “common”) and independent contributions of each predictor (columns in `iv`) to the global *R*^2^ canonical model are computed. In the section below (working examples), we show how the function can conduct the analysis based on blocks of variables as in routine ecological applications of variation partitioning. Response variables (columns in `dv`) must be numerical, while the predictors (`iv`) can be either numerical or categorical. `method` is the type of canonical analysis set as either `RDA` (default), `dbRDA`, or `CCA`. If `method`=“dbRDA”, `dv` should be a distance matrix (i.e., `dist` class). If `dv` is imputed as one numerical vector, RDA would be equivalent to the classic (single response) multiple regression (see Appendix S1). An additional advantage of our package in relation to `relaimpo`, `hier.part` and `dominanceanalysis` is that it also decomposes R^2^adj for multiple regression, so this package is also useful across many other ecological applications and research fields. `type` is the type of total explained variation: “`adjR2`” is for R^2^adj and “`R2`“ for unadjusted *R*^2^, by default is “`adjR2`”`.n.perm` is the number of permutations when computing R^2^adj for CCA and default is 1000 to get a relatively stable value. `trace` is a logical argument indicating whether the output of variation partitioning should be printed (see the example below). It is set by default as FALSE to save screen space, as this output will increase exponentially with the number of predictors. If focusing on variation partitioning, then `trace=TRUE` should be set. `plot.perc` is a logical argument indicating whether a bar plot of the relative independent contribution of predictors is plotted; the default is `FALSE`. The output of `rdacca.hp` is explained in the example below.

## A working example

We illustrate the usage of `rdacca.hp` by using the Doubs River fish data readily available in the `ade4` package (Thioulouse *et al*. 2018). The dataset is a subset of the data originally collected by Verneaux (1973) with the distributions of fish species and environmental factors along the Doubs River in the Jura Mountains, near the France–Switzerland border (also see Verneaux *et al*. 2003). This dataset contains 27 fish species with abundance classes (ranging from 0 to 5) and 11 quantitative environmental variables describing the river morphology and water quality from 30 sites.

We show the results for the variation and hierarchical partitioning of the global R^2^adj of RDA for the Doubs fish data via `rdacca.hp` for three predictors: *alt* (altitude), *oxy* (dissolved oxygen) and *bdo* (biological demand for oxygen) to simplify the presentation of the package rather than provide a complete ecological analysis and related interpretations. These variables (again for presentation convenience) were selected using a stepwise selection based on all environmental predictors using the function `ordistep` in the `vegan` package (Oksanen *et.al*. 2019). Note that the selection of predictors is not a prerequisite of `rdacca.hp` and, as previously noted earlier, can generate biases and incomplete information in variation and hierarchical partitioning. Our function `rdacca.hp` does not limit the number of predictors or groups. Note though, that considering all combinations of all variables is important even though computationally demanding. The selected predictors have a relatively strong correlation structure: a relatively high negative correlation (r=-0.84, *p*<0.001) between *oxy* and *bdo;* a relatively weaker positive correlation between *alt* and *oxy* (r=0.42, *p*=0.02), and a negative correlation between *alt* and *bdo* (r=-0.38, *p*=0.04). This is an important point to make (often missed by practitioners) because the correlation structure in the predictor space (simple pairwise correlations between predictors) may not translate necessarily into high shared fractions among predictors (i.e., semi-partial *R*^2^ calculated on the basis of the predictive space). As such, contrasting the correlation structure among predictors and their correlation structure in predictive space (i.e., variation and hierarchical partitioning) should serve useful in generating insights underlying the relational nature of predictors and responses.

The script below demonstrates the use and outputs of `rdacca.hp` to decompose the global R^2^adj in RDA. Decomposition of R^2^adj in CCA and db-RDA are similar to RDA and shown in the Appendix S2. The standard output of `rdacca.hp` is as follows:

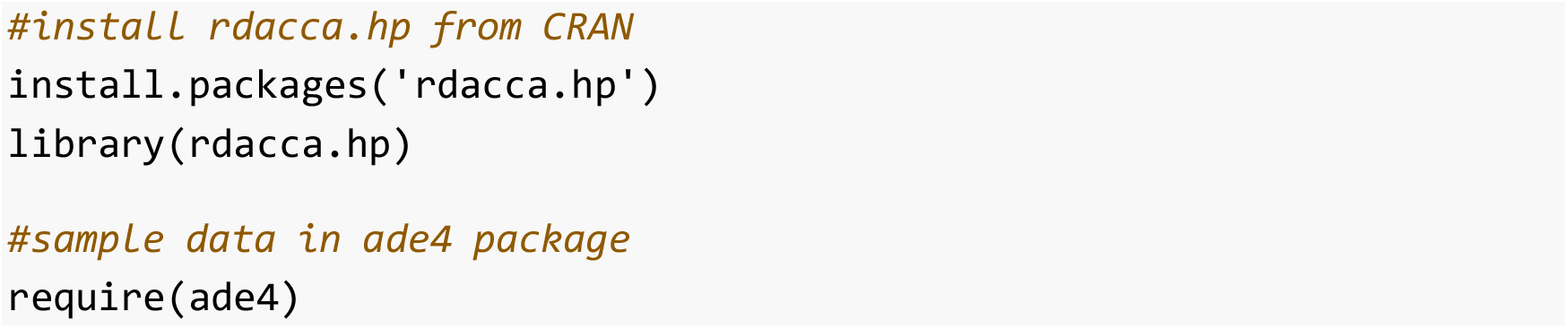
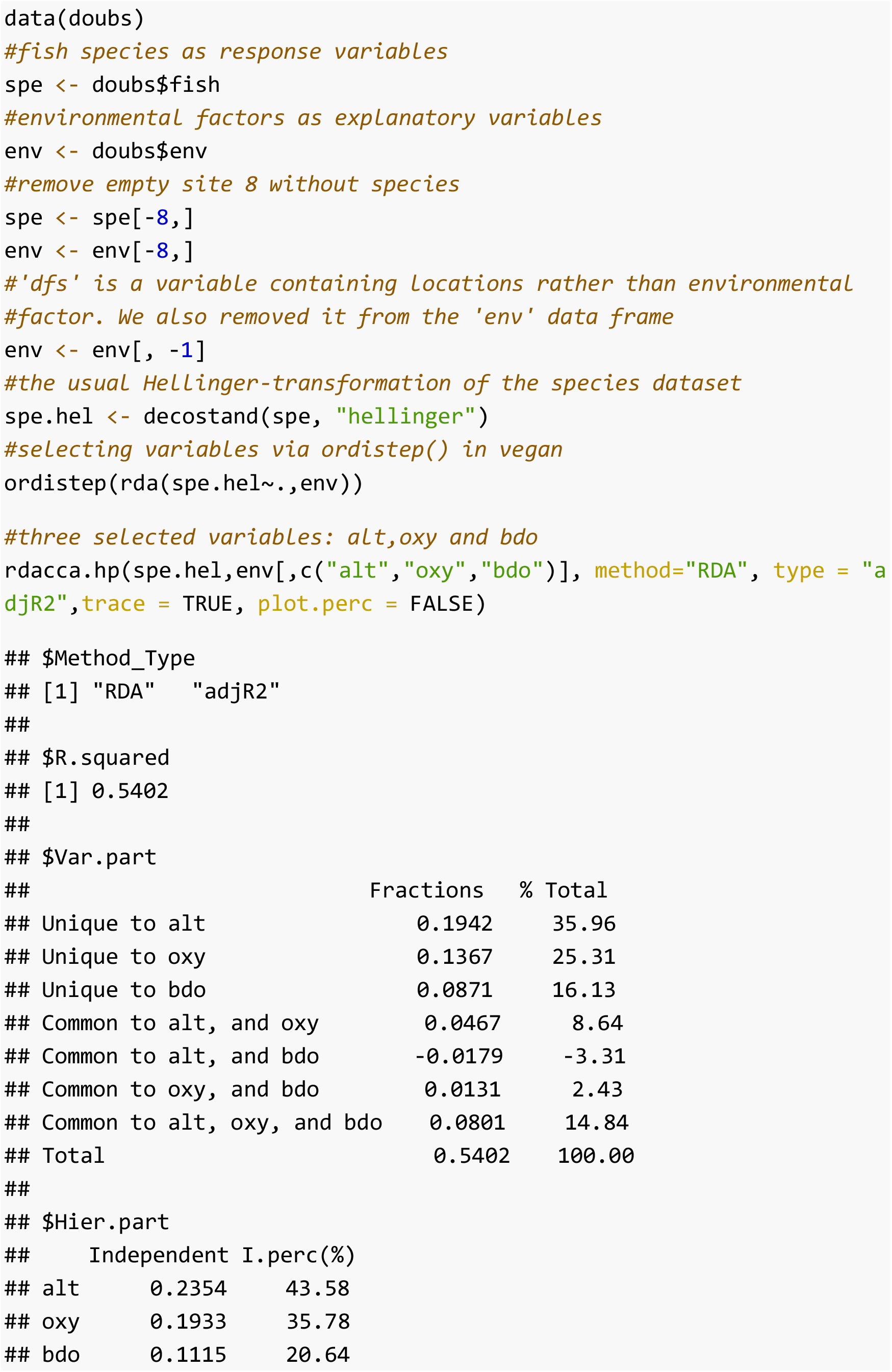

The function `rdacca.hp` returns a list containing three bits (if `trace=FALSE`) or four bits (if `trace=TRUE`).

`$Method_Type:` This bit shows the type of canonical analysis and whether the original or adjusted *R*^2^ were used in the analysis. In the example, it is `RDA` and `adjR2`.

`$R.squared: type=“adjR2”`. In this example, the amount of variation of the fish data matrix explained by the three predictors (**alt**, **oxy** and **bdo**) is 54.02%.

`$Var.part:` It contains output that lists with R^2^adj values for all fractions based on variation partitioning (commonality analysis) if setting `trace=TRUE`. In this example, with three predictors, commonality analysis decomposes R^2^adj into all seven fractions (i.e., 2^3^-1) of unique and common effects. Fractions can be either positive or negative (Nimon & Reio 2011, Ray-Mukherjee *et al* 2014). In this case, the common effect between *alt* and *bod* is negative (−0.0179). Negative common (shared) variation is possible when predictors act as suppressors of other predictors (Pedhazur 1997; Nimon & Reio 2011; Ray-Mukherjee *et a*l 2014).

All fractions (except the residual) sum to the total *R*^2^adj (i.e., the value of `$R.squared`).

`$Hier.part`: This bit is a matrix containing the independent contribution of each predictor in the “Independent” column, and their percentage in the “I.perc” column. In this case, the independent contributions of *alt*, *oxy* and *bdo* are 0.2354, 0.1933 and 0.1115, respectively; note again that they sum to the overall R^2^adj (0.5402). One easily know how to calculate the independent contribution based the result of variation partitioning. For instance, the value (0.1933) for *oxy* is the sum of its unique effect and three average common effects (i.e., 0.1367+ 0.0467/2 +0.0131/2 + 0.0801/3). `rdacca.hp` also produces a bar graph based on the ggplot2 package (Wickham 2016) (Fig. 2). Users are encouraged to use other graph functions to generate other types of plots based on the numerical output of `rdacca.hp`.

**Figure 2.**
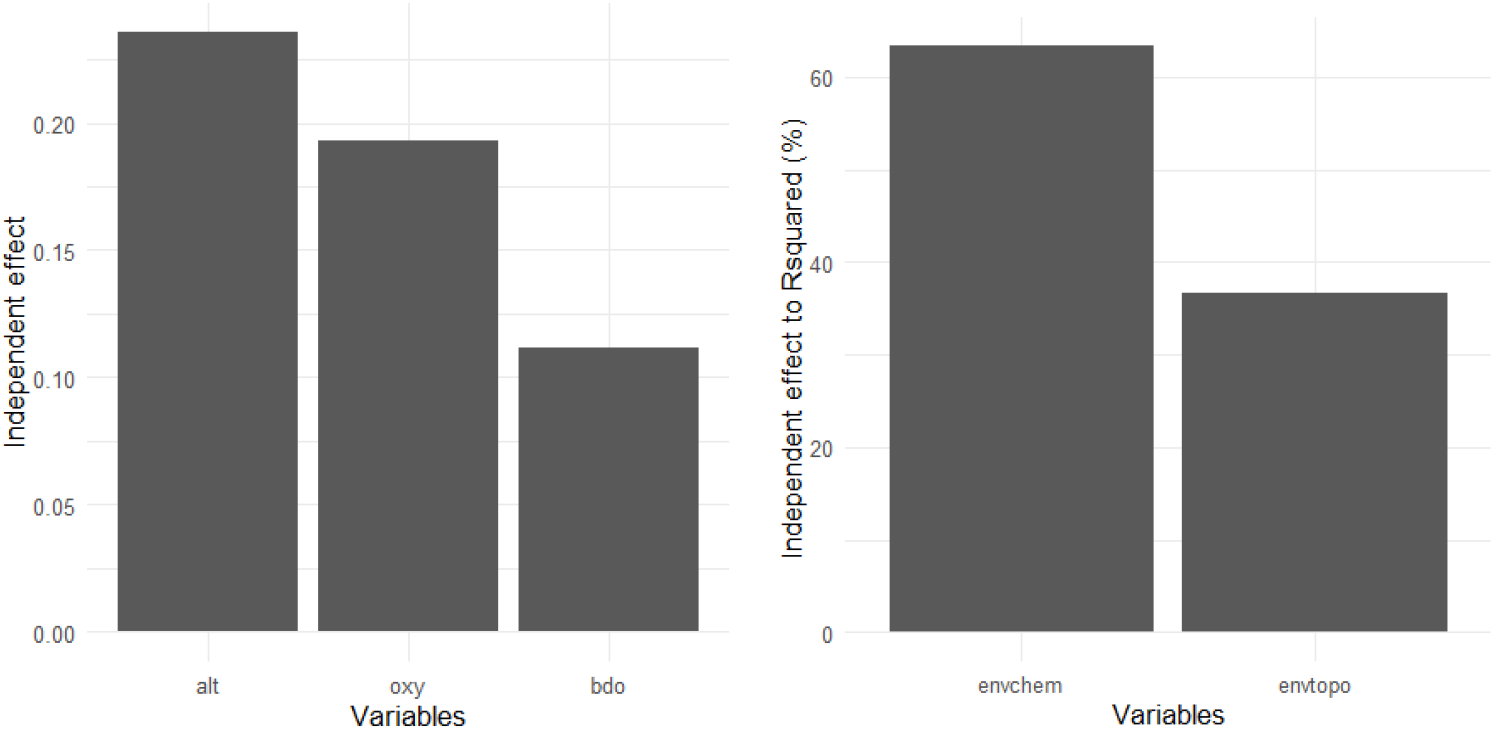
The graphic output of `rdacca.hp` function for the Doubs River fish data. left panel: original independent contribution for three predictors (`plot.perc=FLASE`) and right panel: their relative contribution for two groups to the overall adjusted *R^2^* (`plot.perc=TRUE`).

The following code shows how to apply `rdacca.hp` to conduct a variation partitioning and hierarchical partitioning to two sets of matrices containing environmental variables describing river morphology and water quality. Note `rdacca.hp` has no limit for the number of sets of predictor matrices and an example with five predictor groups is provided in the Appendix S3. rdacca. hp does not include the traditional representation of variation partitioning as a Venn diagram, which becomes impossible for more than four predictors.

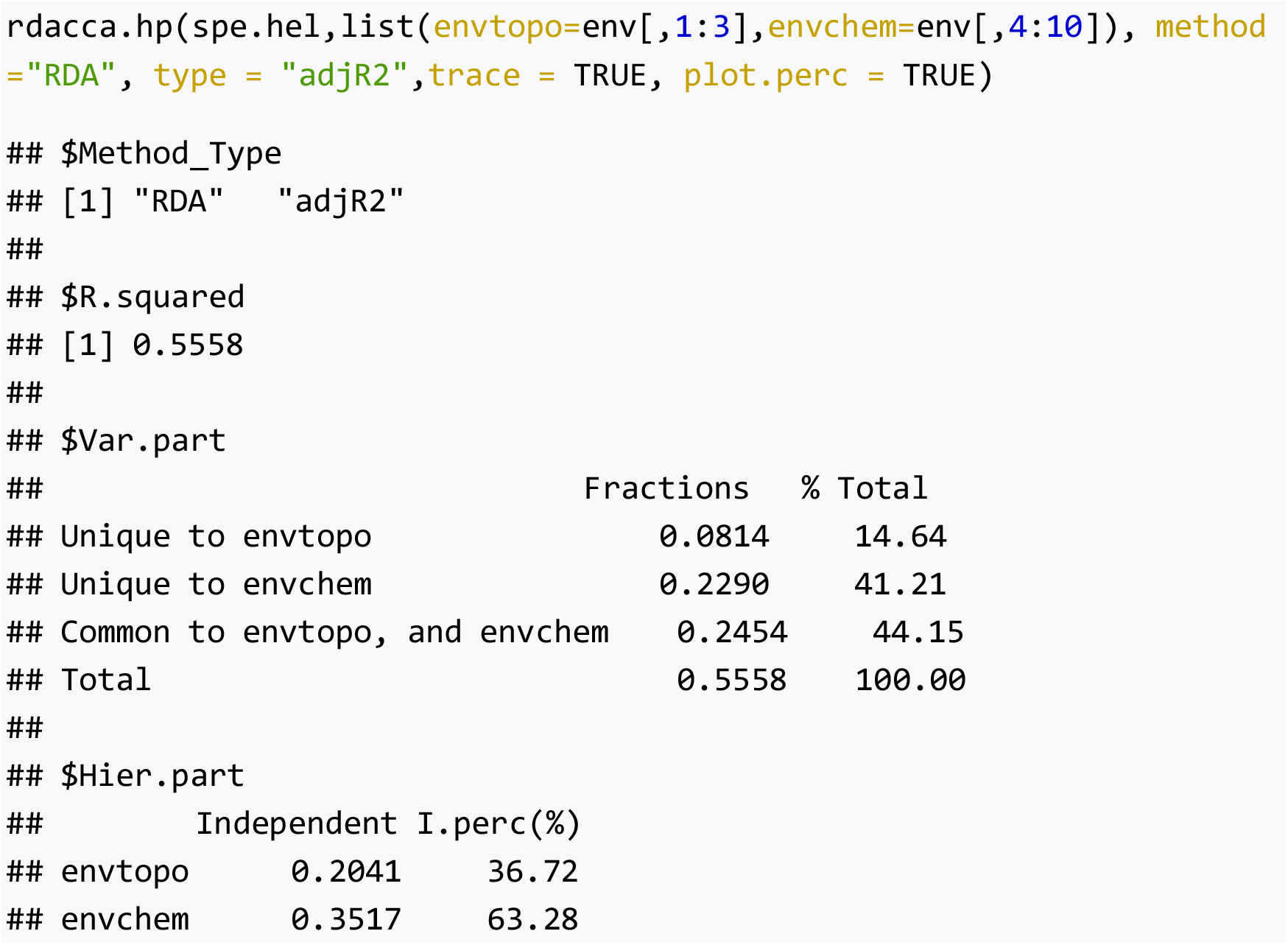

## Discussion

Multivariate regressions and canonical analysis are quintessential quantitative frameworks to tackle many ecological and evolutionary problems. Variation partitioning and hierarchical partitioning (HP) are frameworks that allow going beyond the usual and standard interpretation of partial slopes (see Ray-Mukherjee *et al*. 2014 for a review). That said, our presentation of HP allows realizing that both methods are interrelated, and that variation partitioning can serve as a precursor of HP. While variation partitioning estimates unique and common variation in a single full model containing all variables, HP is based on principles of all subset regression and model averaging which are known to improve model inference and interpretability over traditional model selection procedures (Burnham & Anderson 2002). The two main motivations for using variation partitioning and HP over traditional model selection procedures are that: a) candidate predictors are conditional on the variance explained by the models already retained by the selection procedure (Thompson 1995; Nathans, Oswald & Nimon 2012). As such, a predictor that can be quite relevant in multiple sub-models (indicating overall importance) may not be retained by model selection procedures; b) model selection can be heavily influenced by sampling variation (Thompson 1995; Nathans, Oswald & Nimon 2012); if another sample were to be used, the selected predictors could vary. By using HP, one can analyze variable importance over all possible models.

While most canonical analysis is used to analyze species distributional matrices, our package can also motivate ecologists to explore its use to different types of response matrices and problems (e.g., multiple traits). Note that we decided not to consider significance testing for independent contributions in HP because predictor (or groups of predictors) relative importance is usually considered as an exploratory framework for interpreting regression rather than an inferential tool. Finally, one disadvantage of the variation partitioning and HP frameworks is that the calculation volume increases exponentially with the increase of predictor variables. We will continue to optimize our package to improve, among other issues, the calculation speed. Although we demonstrated the application of `rdacca.hp` package using an ecology example, this package is also applicable to any other areas where canonical analysis and multiple regressions are applied. Currently, our `rdacca.hp` package has been used in peer-reviewed papers (e.g. Li *et al*. 2020; Song *et al*. 2020; Sun *et al*. 2020; Wang *et al*. 2020; Xiong *et al*. 2020; Zhou *et al*. 2020). Researchers using the `rdacca.hp` package in their studies, should cite this article and the rdacca.hp package as well. Citation information can be obtained by typing: `citation(“rdacca.hp”`).

## Acknowledgements

The research was supported by the Strategic Priority Research Program of the Chinese Academy of Sciences (XDA19050404) and the National Science and Technology Basic Resources Survey Program of China (2019FY100204). PP-N was supported by the Canada Research Chair (CRC) program. Otherwise there is no conflict of interest among the authors.

## Authors’ contributions

JSL conceived the idea. JSL and PP-N wrote the package and conducted the analysis. All authors participated in writing multiples drafts.

