## appendix S1 for "rdacca.hp: an R package for generalizing hierarchical and variation partitioning in multiple regression and canonical analysis"

```
#We show that the results from package relaimpo, heir.part and dominanc#-eanalysis are identical for multiple regression based on same data
#install.packages("hier.part")
library(hier.part)
#example data: urbanwq in hier.part package
data(urbanwq)
env <- urbanwq[,2:8]
#the goodness-of-fit choice for heir.part has to be chosen as R2 (gof= #Rsqu)
hier.part(urbanwq$lec, env, fam = "gaussian", gof = "Rsqu")
```

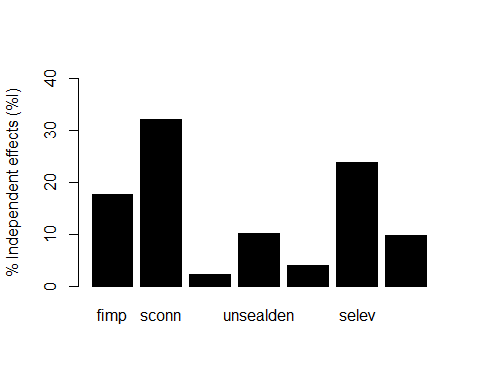

```
## $gfs
##   [1] 0.00000000 0.59047318 0.82682075 0.01098174 0.39635905 0.12282709
##   [7] 0.62445471 0.33033972 0.83410378 0.59758915 0.62061433 0.65284362
##  [13] 0.77159448 0.60702109 0.82683793 0.82691037 0.83582029 0.85403927
##  [19] 0.84727070 0.41798147 0.13980912 0.76026824 0.33080645 0.39744819
##  [25] 0.66503431 0.52650125 0.67608730 0.43330271 0.64633406 0.83447868
##  [31] 0.83411192 0.83965414 0.85551898 0.84727151 0.62061622 0.65949970
##  [37] 0.82672399 0.62672410 0.65445206 0.77226833 0.62512912 0.77811174
##  [43] 0.66312305 0.82026934 0.82691040 0.84141253 0.87751308 0.84785650
##  [49] 0.83932285 0.85513956 0.84985538 0.85490360 0.85107746 0.87685251
##  [55] 0.43348171 0.81801707 0.55827372 0.76851361 0.50536622 0.77107209
##  [61] 0.73969961 0.53654123 0.68316625 0.67713148 0.83451931 0.84541929
##  [67] 0.87865868 0.84793008 0.84176310 0.85633390 0.85010056 0.85662169
##  [73] 0.85116200 0.88236321 0.66297682 0.84039075 0.62908864 0.82701685
##  [79] 0.66414800 0.85116957 0.78348135 0.66312339 0.82956916 0.85200153
##  [85] 0.84338543 0.87900342 0.85317612 0.87755498 0.85162358 0.89200389
##  [91] 0.85535421 0.85816222 0.88318976 0.88611089 0.83628377 0.61132790
##  [97] 0.82521005 0.77285776 0.74013871 0.84631087 0.88052535 0.85779318
## [103] 0.87866729 0.85162457 0.89525987 0.85679323 0.86058475 0.89708584
## [109] 0.89557323 0.84575908 0.66461591 0.85137732 0.86367597 0.85322499
## [115] 0.87916623 0.85818567 0.89224695 0.89515882 0.88779139 0.83642968
## [121] 0.88093891 0.86223879 0.89900950 0.90101494 0.90231612 0.86447417
## [127] 0.89516613 0.90324171
##
## $IJ
##                    I            J      Total
## fimp      0.16018583  0.430287348 0.59047318
## sconn     0.28970489  0.537115864 0.82682075
## sdensep   0.02090086 -0.009919125 0.01098174
## unsealden 0.09181298  0.304546072 0.39635905
## fcarea    0.03609565  0.086731439 0.12282709
## selev     0.21589289  0.408561821 0.62445471
## amgeast   0.08864861  0.241691110 0.33033972
##
## $I.perc
##           ind.exp.var
## fimp        17.734548
## sconn       32.073905
## sdensep      2.313983
## unsealden   10.164829
## fcarea       3.996234
## selev       23.902006
## amgeast      9.814495
##
## $params
## $params$full.model
## [1] "y ~ fimp + sconn + sdensep + unsealden + fcarea + selev + amgeast"
##
## $params$family
## [1] "gaussian"
##
## $params$link
## [1] "default"
##
## $params$gof
## [1] "Rsqu"
lm.1=lm(urbanwq$lec~.,env)
#install.packages("relaimpo")
library(relaimpo)
## Loading required package: MASS
## Loading required package: boot
## Loading required package: survey
## Loading required package: grid
## Loading required package: Matrix
## Loading required package: survival
##
## Attaching package: 'survival'
## The following object is masked from 'package:boot':
##
##     aml
##
## Attaching package: 'survey'
## The following object is masked from 'package:graphics':
##
##     dotchart
## Loading required package: mitools
## This is the global version of package relaimpo.
## If you are a non-US user, a version with the interesting additional metric pmvd is available
## from Ulrike Groempings web site at prof.beuth-hochschule.de/groemping.
calc.relimp(lm.1)
## Response variable: urbanwq$lec
## Total response variance: 0.1239396
## Analysis based on 15 observations
##
## 7 Regressors:
## fimp sconn sdensep unsealden fcarea selev amgeast
## Proportion of variance explained by model: 90.32%
## Metrics are not normalized (rela=FALSE).
##
## Relative importance metrics:
##
##                  lmg
## fimp      0.16018583
## sconn     0.28970489
## sdensep   0.02090086
## unsealden 0.09181298
## fcarea    0.03609565
## selev     0.21589289
## amgeast   0.08864861
##
## Average coefficients for different model sizes:
##
##                      1X           2Xs           3Xs           4Xs           5Xs
## fimp       1.438029e+00  1.051726e+00  7.663988e-01  5.796452e-01  4.885536e-01
## sconn      8.608541e-01  8.792503e-01  8.798651e-01  8.522788e-01  7.848919e-01
## sdensep   -9.625479e-03  1.068006e-02  1.588652e-02  1.612028e-02  1.429844e-02
## unsealden -3.641278e+01 -2.163258e+01 -1.055358e+01 -3.143680e+00  6.885511e-01
## fcarea     4.822311e-01  1.962823e-01  9.951369e-02  3.054207e-02 -4.400250e-02
## selev     -8.760166e-02 -7.589446e-02 -6.784185e-02 -6.406297e-02 -6.297468e-02
## amgeast   -2.151702e-05 -7.864678e-06  9.207176e-07  6.917650e-06  1.075835e-05
##                     6Xs           7Xs
## fimp       5.161968e-01  6.263840e-01
## sconn      6.929782e-01  6.084176e-01
## sdensep    1.055266e-02  5.349108e-03
## unsealden  3.372312e+00  6.055790e+00
## fcarea    -1.146659e-01 -1.624194e-01
## selev     -6.212648e-02 -5.896531e-02
## amgeast    1.356463e-05  1.597516e-05
#install.packages('dominanceanalysis')
library(dominanceanalysis)
dominanceAnalysis(lm.1)
##
## * Fit index:  r2
##
## Average contribution of each variable:
##
##     sconn     selev      fimp unsealden   amgeast    fcarea  sdensep
##     0.290     0.216     0.160     0.092     0.089     0.036    0.021
#rdacca.hp can work for multiple regression, its result is identical to#former packages
#install.packages('rdacca.hp')
library(rdacca.hp)
## Loading required package: vegan
## Loading required package: permute
## Loading required package: lattice
##
## Attaching package: 'lattice'
## The following object is masked from 'package:boot':
##
##     melanoma
## This is vegan 2.5-6
##
## Attaching package: 'vegan'
## The following object is masked from 'package:survey':
##
##     calibrate
## Loading required package: ggplot2
rdacca.hp(urbanwq$lec, env,method="RDA", type = "R2")
```

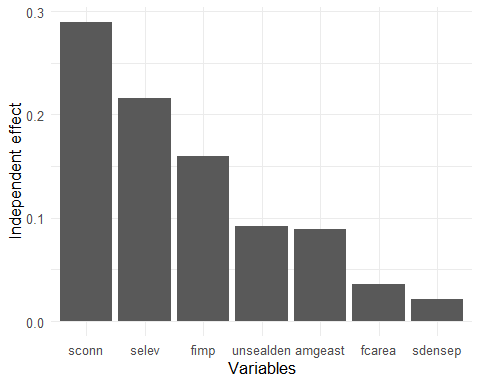

```
## $Method_Type
## [1] "RDA" "R2"
##
## $R.squared
## [1] 0.9032
##
## $Hier.part
##           Independent I.perc(%)
## fimp           0.1602     17.74
## sconn          0.2897     32.07
## sdensep        0.0209      2.31
## unsealden      0.0918     10.16
## fcarea         0.0361      4.00
## selev          0.2159     23.90
## amgeast        0.0886      9.81
#An additional advantage of rdacca.hp in relation to former packages is#that it also decomposes adjusted #Rsquared for multiple regression
rdacca.hp(urbanwq$lec, env,method="RDA",type = "adjR2")
```

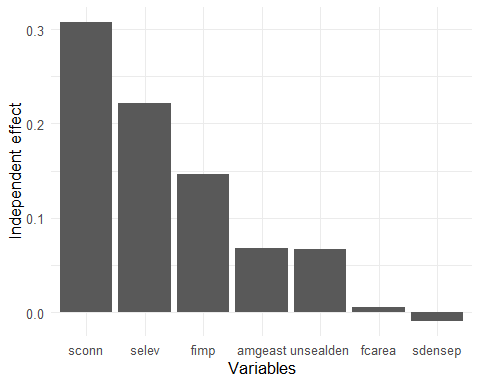

```
## $Method_Type
## [1] "RDA"   "adjR2"
##
## $R.squared
## [1] 0.8066
##
## $Hier.part
##           Independent I.perc(%)
## fimp           0.1461     18.11
## sconn          0.3078     38.16
## sdensep       -0.0093     -1.15
## unsealden      0.0665      8.24
## fcarea         0.0059      0.73
## selev          0.2220     27.52
## amgeast        0.0676      8.38
```
