## appendix S2 for "rdacca.hp: an R package for generalizing hierarchical and variation partitioning in multiple regression and canonical analysis"

```
#install rdacca.hp from CRAN or github and load it
# install.packages('rdacca.hp')
library(rdacca.hp)
## Loading required package: vegan
## Loading required package: permute
## Loading required package: lattice
## This is vegan 2.5-7
## Loading required package: ggplot2
#library(devtools)
#install_github('laijiangshan/rdacca.hp')
#sample data in ade4 package
require(ade4)
## Loading required package: ade4
data(doubs)
# Fish species as response variables
spe <- doubs$fish
# Environmental factors as explanatory variables
env <- doubs$env
#Remove empty site 8 without species
spe <- spe[-8,]
env <- env[-8,]
# Apart from being a spatial position varialbe rather than
# environmental variable, remove the 'dfs' variable (the distance
# from the source) from the 'env' data frame
env <- env[, -1]
# Hellinger-transformation of the species dataset for RDA to deal # with the 'double zero' problem
spe.hel <- decostand(spe, "hellinger")
#variables selection for RDA via ordistep() in vegan package
ordistep(rda(spe.hel~.,env))
##
## Start: spe.hel ~ alt + slo + flo + pH + har + pho + nit + amm + oxy +      bdo
##
##       Df     AIC      F Pr(>F)
## - amm  1 -36.675 0.4025  0.840
## - nit  1 -36.556 0.4781  0.810
## - bdo  1 -36.361 0.6028  0.660
## - pho  1 -36.129 0.7525  0.600
## - pH   1 -36.328 0.6241  0.575
## - alt  1 -35.712 1.0240  0.370
## - slo  1 -35.043 1.4682  0.160
## - flo  1 -34.355 1.9354  0.130
## - har  1 -34.405 1.9012  0.095 .
## - oxy  1 -32.507 3.2469  0.030 *
## ---
## Signif. codes:  0 '***' 0.001 '**' 0.01 '*' 0.05 '.' 0.1 ' ' 1
##
## Step: spe.hel ~ alt + slo + flo + pH + har + pho + nit + oxy + bdo
##
##       Df     AIC      F Pr(>F)
## - nit  1 -37.744 0.6202  0.650
## - pH   1 -37.682 0.6617  0.620
## - pho  1 -37.420 0.8406  0.435
## - bdo  1 -37.209 0.9850  0.380
## - alt  1 -37.074 1.0785  0.290
## - har  1 -35.788 1.9893  0.125
## - slo  1 -36.449 1.5161  0.120
## - flo  1 -35.775 1.9981  0.090 .
## - oxy  1 -33.601 3.6328  0.015 *
## ---
## Signif. codes:  0 '***' 0.001 '**' 0.01 '*' 0.05 '.' 0.1 ' ' 1
##
## Step: spe.hel ~ alt + slo + flo + pH + har + pho + oxy + bdo
##
##       Df     AIC      F Pr(>F)
## - pH   1 -38.798 0.6632  0.565
## - pho  1 -38.450 0.9127  0.440
## - slo  1 -37.553 1.5692  0.150
## - bdo  1 -37.121 1.8929  0.105
## - flo  1 -36.914 2.0501  0.045 *
## - har  1 -36.319 2.5069  0.045 *
## - alt  1 -34.788 3.7271  0.020 *
## - oxy  1 -32.076 6.0530  0.005 **
## ---
## Signif. codes:  0 '***' 0.001 '**' 0.01 '*' 0.05 '.' 0.1 ' ' 1
##
## Step: spe.hel ~ alt + slo + flo + har + pho + oxy + bdo
##
##       Df     AIC      F Pr(>F)
## - pho  1 -39.542 0.9294  0.450
## - slo  1 -39.010 1.3351  0.205
## - flo  1 -38.115 2.0352  0.145
## - bdo  1 -38.223 1.9496  0.120
## - har  1 -37.487 2.5399  0.020 *
## - alt  1 -35.825 3.9278  0.020 *
## - oxy  1 -32.358 7.0935  0.005 **
## ---
## Signif. codes:  0 '***' 0.001 '**' 0.01 '*' 0.05 '.' 0.1 ' ' 1
##
## Step: spe.hel ~ alt + slo + flo + har + oxy + bdo
##
##       Df     AIC      F Pr(>F)
## - slo  1 -39.807 1.3559  0.230
## - har  1 -38.393 2.5232  0.070 .
## - flo  1 -38.891 2.1054  0.050 *
## - alt  1 -36.648 4.0443  0.005 **
## - bdo  1 -35.341 5.2450  0.005 **
## - oxy  1 -32.796 7.7443  0.005 **
## ---
## Signif. codes:  0 '***' 0.001 '**' 0.01 '*' 0.05 '.' 0.1 ' ' 1
##
## Step: spe.hel ~ alt + flo + har + oxy + bdo
##
##       Df     AIC      F Pr(>F)
## - flo  1 -39.394 1.9959  0.115
## - har  1 -38.436 2.8358  0.045 *
## - alt  1 -36.339 4.7732  0.015 *
## - bdo  1 -35.290 5.7958  0.010 **
## - oxy  1 -32.529 8.6721  0.005 **
## ---
## Signif. codes:  0 '***' 0.001 '**' 0.01 '*' 0.05 '.' 0.1 ' ' 1
##
## Step: spe.hel ~ alt + har + oxy + bdo
##
##       Df     AIC      F Pr(>F)
## - har  1 -38.789 2.2555  0.060 .
## - bdo  1 -34.740 6.1895  0.010 **
## - alt  1 -31.966 9.2198  0.005 **
## - oxy  1 -31.884 9.3140  0.005 **
## ---
## Signif. codes:  0 '***' 0.001 '**' 0.01 '*' 0.05 '.' 0.1 ' ' 1
## Call: rda(formula = spe.hel ~ alt + har + oxy + bdo, data = env)
##
##               Inertia Proportion Rank
## Total          0.5025     1.0000
## Constrained    0.3139     0.6247    4
## Unconstrained  0.1886     0.3753   24
## Inertia is variance
##
## Eigenvalues for constrained axes:
##    RDA1    RDA2    RDA3    RDA4
## 0.22284 0.05131 0.02781 0.01197
##
## Eigenvalues for unconstrained axes:
##     PC1     PC2     PC3     PC4     PC5     PC6     PC7     PC8
## 0.04919 0.03983 0.02566 0.01708 0.01273 0.01156 0.00757 0.00594
## (Showing 8 of 24 unconstrained eigenvalues)
# three selected variables: alt,oxy and bdo
rdacca.hp(spe.hel,env[,c("alt","oxy","bdo")], method="RDA", type = "adjR2",trace = TRUE)
```

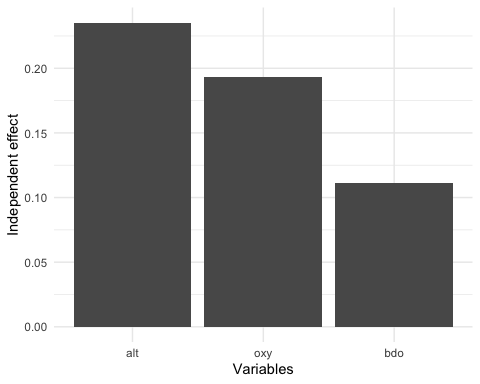

```
## $Method_Type
## [1] "RDA"   "adjR2"
##
## $R.squared
## [1] 0.5402
##
## $Var.part
##                             Fractions   % Total
## Unique to alt                  0.1942     35.96
## Unique to oxy                  0.1367     25.31
## Unique to bdo                  0.0871     16.13
## Common to alt, and oxy         0.0467      8.64
## Common to alt, and bdo        -0.0179     -3.31
## Common to oxy, and bdo         0.0131      2.43
## Common to alt, oxy, and bdo    0.0801     14.84
## Total                          0.5402    100.00
##
## $Hier.part
##     Independent I.perc(%)
## alt      0.2354     43.58
## oxy      0.1933     35.78
## bdo      0.1115     20.64
# conduct a variation portioning and hierarchical partitioning to two sets of# matrices containing environmental variables describing river morphology and# water quality
rdacca.hp(spe.hel,list(envtopo=env[,1:3],envchem=env[,4:10]), method="RDA", type = "adjR2",trace = TRUE)
```

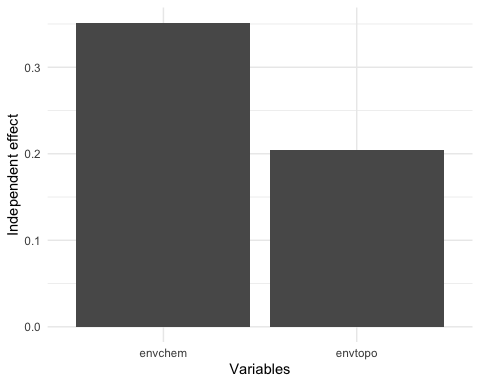

```
## $Method_Type
## [1] "RDA"   "adjR2"
##
## $R.squared
## [1] 0.5558
##
## $Var.part
##                                Fractions   % Total
## Unique to envtopo                 0.0814     14.64
## Unique to envchem                 0.2290     41.21
## Common to envtopo, and envchem    0.2454     44.15
## Total                             0.5558    100.00
##
## $Hier.part
##         Independent I.perc(%)
## envtopo      0.2041     36.72
## envchem      0.3517     63.28
#example for CCA
rdacca.hp(spe,env[,c("alt","oxy","bdo")], method="CCA", type = "adjR2",trace = TRUE)
```

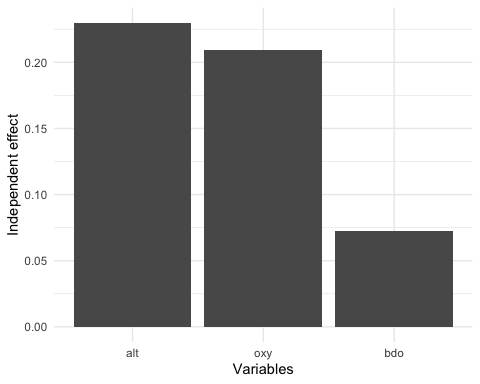

```
## $Method_Type
## [1] "CCA"   "adjR2"
##
## $R.squared
## [1] 0.5121
##
## $Var.part
##                             Fractions   % Total
## Unique to alt                  0.1724     33.67
## Unique to oxy                  0.1532     29.92
## Unique to bdo                  0.0553     10.80
## Common to alt, and oxy         0.0871     17.00
## Common to alt, and bdo         0.0088      1.73
## Common to oxy, and bdo         0.0062      1.21
## Common to alt, oxy, and bdo    0.0291      5.68
## Total                          0.5121    100.00
##
## $Hier.part
##     Independent I.perc(%)
## alt      0.2301     44.93
## oxy      0.2095     40.91
## bdo      0.0725     14.16
#example for db-RDA
rdacca.hp(vegdist(spe),env[,c("alt","oxy","bdo")], method="dbRDA", type = "adjR2",trace = TRUE)
```

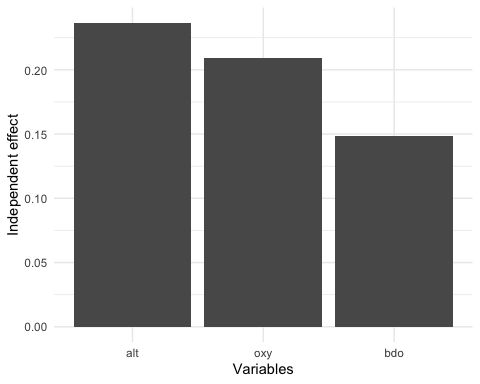

```
## $Method_Type
## [1] "dbRDA" "adjR2"
##
## $R.squared
## [1] 0.5948
##
## $Var.part
##                             Fractions   % Total
## Unique to alt                  0.1939     32.59
## Unique to oxy                  0.1664     27.98
## Unique to bdo                  0.1468     24.69
## Common to alt, and oxy         0.0597     10.04
## Common to alt, and bdo        -0.0232     -3.90
## Common to oxy, and bdo        -0.0229     -3.85
## Common to alt, oxy, and bdo    0.0741     12.45
## Total                          0.5948    100.00
##
## $Hier.part
##     Independent I.perc(%)
## alt      0.2368     39.81
## oxy      0.2095     35.22
## bdo      0.1485     24.97
```

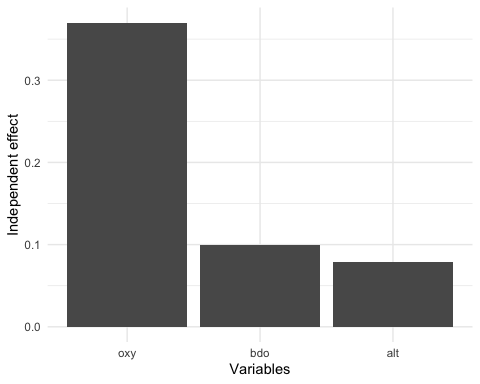

```
## $Method_Type
## [1] "RDA" "R2"
##
## $R.squared
## [1] 0.5493
##
## $Hier.part
##     Independent I.perc(%)
## alt      0.0790     14.38
## oxy      0.3702     67.39
## bdo      0.1001     18.22
```
