## appendix S3 for "rdacca.hp: an R package for generalizing hierarchical and variation partitioning in multiple regression and canonical analysis"

```
#show rdacca.hp for mite data with categorical predictors.
#install.packages('rdacca.hp')
library(rdacca.hp)
## Loading required package: vegan
## Loading required package: permute
## Loading required package: lattice
## This is vegan 2.5-6
## Loading required package: ggplot2
library(vegan)
data(mite)
data(mite.env)
#Hellinger-transform the species dataset
mite.hel <- decostand(mite, "hellinger")
rdacca.hp(mite.hel,mite.env,method="RDA",type="adjR2")
```

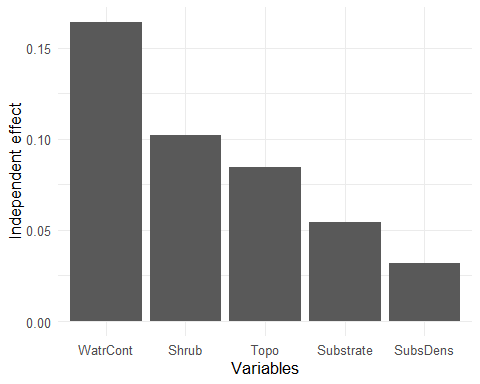

```
## $Method_Type
## [1] "RDA"   "adjR2"
##
## $R.squared
## [1] 0.4366
##
## $Hier.part
##           Independent I.perc(%)
## SubsDens       0.0320      7.33
## WatrCont       0.1642     37.61
## Substrate      0.0543     12.44
## Shrub          0.1018     23.32
## Topo           0.0843     19.31
rdacca.hp(vegdist(mite),mite.env,method="dbRDA",type="adjR2")
```

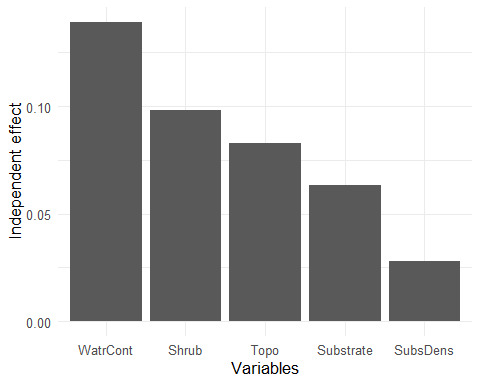

```
## $Method_Type
## [1] "dbRDA" "adjR2"
##
## $R.squared
## [1] 0.4113
##
## $Hier.part
##           Independent I.perc(%)
## SubsDens       0.0280      6.81
## WatrCont       0.1393     33.87
## Substrate      0.0631     15.34
## Shrub          0.0980     23.83
## Topo           0.0829     20.16
rdacca.hp(mite,mite.env,method="CCA",type="adjR2")
```

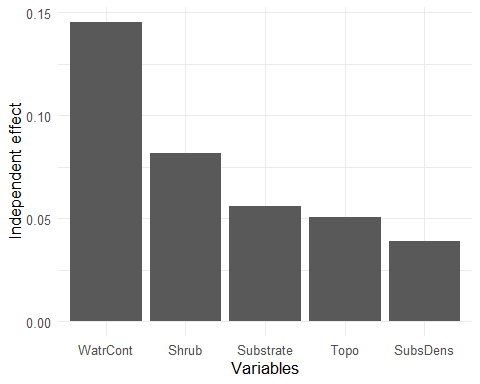

```
## $Method_Type
## [1] "CCA"   "adjR2"
##
## $R.squared
## [1] 0.3721
##
## $Hier.part
##           Independent I.perc(%)
## SubsDens       0.0388     10.43
## WatrCont       0.1453     39.05
## Substrate      0.0558     15.00
## Shrub          0.0817     21.96
## Topo           0.0505     13.57
#conduct a variation portioning and hierarchical partitioning for five sets #of matrices
data(mite.xy)
data(mite.pcnm)
iv <- list(env1=mite.env[,1:2],env2=mite.env[,3:5],xy=mite.xy,pcnm1=mite.pcnm[,1:7],pcnm2=mite.pcnm[,8:22])
rdacca.hp(mite.hel,iv,method="RDA",trace = TRUE,plot.perc = FALSE)
```

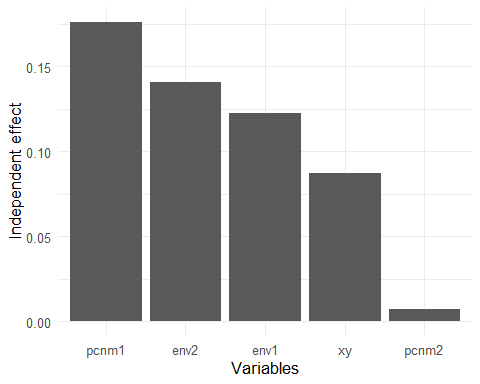

```
## $Method_Type
## [1] "RDA"   "adjR2"
##
## $R.squared
## [1] 0.5335
##
## $Var.part
##                                            Fractions   % Total
## Unique to env1                                0.0318      5.96
## Unique to env2                                0.0334      6.25
## Unique to xy                                 -0.0044     -0.83
## Unique to pcnm1                               0.0524      9.82
## Unique to pcnm2                               0.0010      0.18
## Common to env1, and env2                      0.0131      2.45
## Common to env1, and xy                        0.0073      1.37
## Common to env2, and xy                        0.0049      0.92
## Common to env1, and pcnm1                     0.0180      3.37
## Common to env2, and pcnm1                     0.1071     20.07
## Common to xy, and pcnm1                       0.0359      6.73
## Common to env1, and pcnm2                     0.0206      3.86
## Common to env2, and pcnm2                     0.0066      1.24
## Common to xy, and pcnm2                       0.0283      5.30
## Common to pcnm1, and pcnm2                    0.0093      1.74
## Common to env1, env2, and xy                  0.0010      0.18
## Common to env1, env2, and pcnm1               0.0192      3.60
## Common to env1, xy, and pcnm1                 0.0198      3.71
## Common to env2, xy, and pcnm1                 0.0300      5.62
## Common to env1, env2, and pcnm2               0.0068      1.27
## Common to env1, xy, and pcnm2                 0.0190      3.57
## Common to env2, xy, and pcnm2                -0.0084     -1.58
## Common to env1, pcnm1, and pcnm2             -0.0126     -2.36
## Common to env2, pcnm1, and pcnm2             -0.0366     -6.87
## Common to xy, pcnm1, and pcnm2               -0.0256     -4.79
## Common to env1, env2, xy, and pcnm1           0.1970     36.93
## Common to env1, env2, xy, and pcnm2           0.0056      1.05
## Common to env1, env2, pcnm1, and pcnm2        0.0011      0.20
## Common to env1, xy, pcnm1, and pcnm2          0.0183      3.42
## Common to env2, xy, pcnm1, and pcnm2         -0.0068     -1.28
## Common to env1, env2, xy, pcnm1, and pcnm2   -0.0593    -11.11
## Total                                         0.5335    100.00
##
## $Hier.part
##       Independent I.perc(%)
## env1       0.1227     23.00
## env2       0.1405     26.34
## xy         0.0873     16.36
## pcnm1      0.1761     33.01
## pcnm2      0.0069      1.29
```
